# SRC Knockdown Impairs Proliferation, Migration, and Invasion While Promoting Apoptosis in HTR8/SVneo Trophoblast Cells via Activation of the PI3K/Akt/Bcl-2 Signaling Pathway

**DOI:** 10.64898/2026.05.28.728375

**Authors:** Yao He, Hai-Tao Pan, Guo-Ping Li, Feng Zhang, Ye-Jun Jiang, Gui-Yu Xia, Jian Zhao, Jin-Long Ding, Xin-Yue Zhang, Na Ding, Hai-Gang Ding, Bin Yu

## Abstract

SRC knockdown inhibits trophoblast cell proliferation, migration, and invasion while inducing apoptosis via activation of the PI3K/Akt/Bcl-2 signaling pathway. Trophoblast dysfunction is central to pregnancy disorders such as preeclampsia and miscarriage, yet the role of SRC, a non-receptor tyrosine kinase, in these cells remains poorly understood. This study aimed to elucidate the functional impact of SRC on trophoblast behavior and its underlying mechanism. Using siRNA-mediated knockdown in HTR8/SVneo cells, we confirmed efficient reduction of SRC mRNA and protein expression via RT-qPCR and Western blot. Functional assays demonstrated that SRC silencing significantly suppressed cell proliferation (CCK-8), migration (wound healing), and invasion (Transwell), while promoting apoptosis, evidenced by increased Annexin V-FITC/PI staining and upregulated Caspase-3 and Caspase-9 protein levels. Mechanistically, Western blot analysis revealed that SRC knockdown upregulated PI3K, Akt1, and Bcl-2 protein expression without altering IRS1 levels, indicating activation of the PI3K/Akt/Bcl-2 pro-survival pathway. This paradoxical activation appears to be a compensatory feedback insufficient to overcome SRC loss-induced dysfunction. Our findings identify SRC as a critical positive regulator of trophoblast proliferation, motility, and survival, acting through a non-canonical, IRS1-independent negative regulation of PI3K/Akt signaling. This study provides novel insights into trophoblast biology and suggests SRC as a potential therapeutic target for pregnancy complications; future in vivo studies are warranted to validate these mechanisms.

## 1. Introduction

Proper placentation is a fundamental prerequisite for a successful pregnancy, as it establishes the critical interface for nutrient, gas, and waste exchange between the mother and the developing fetus. The functional integrity of the placenta is primarily governed by the trophoblast cells, a heterogeneous population that originates from the outer layer of the blastocyst. In particular, extravillous trophoblasts (EVTs) undergo a precisely regulated program of proliferation, migration, and invasion into the maternal decidua, a process essential for remodeling the spiral arteries and ensuring adequate uteroplacental blood flow. Dysregulation of these intricate trophoblast functions is a recognized etiological factor for a spectrum of pregnancy complications, including early pregnancy loss, preeclampsia (PE), and intrauterine growth restriction (FGR) [1]. These disorders represent a significant burden in maternal-fetal medicine, with preeclampsia affecting approximately 5-8% of all pregnancies globally and remaining a leading cause of maternal and perinatal morbidity and mortality. Despite this clinical importance, the detailed molecular mechanisms that orchestrate EVT biology, particularly the signaling networks controlling their survival, motility, and invasive capacity, remain incompletely understood.

The Src family of non-receptor tyrosine kinases, of which the proto-oncogene Src is the founding member, plays a pivotal role in transducing signals from various cell surface receptors to regulate a multitude of cellular processes, including proliferation, adhesion, migration, and survival. While the role of Src has been extensively characterized in the context of cancer progression and metastasis, its specific functions within the placenta are less well-defined. Related research has demonstrated that signaling molecules such as Shp2, a protein tyrosine phosphatase, critically regulates trophoblast cell cycle progression through modulation of the p53-p21 pathway and the PI3K-Akt and MAPK signaling cascades [2]. Furthermore, steroid receptor coactivator-2 (SRC-2) has been shown to be essential for the viability, migration, and invasion of human EVTs [1]. However, the specific function of the prototypical Src itself, particularly its role in integrating downstream signaling to control trophoblast cell fate and function, is a significant knowledge gap. The potential for Src to act as a central hub, simultaneously influencing multiple key cellular behaviors in trophoblasts, warrants focused investigation. Thus, elucidating the specific molecular pathways through which Src governs trophoblast function is crucial for advancing our understanding of normal placental development and the pathogenesis of placental disorders.

The PI3K/Akt signaling pathway is a well-established and central regulator of cell survival, proliferation, and metabolism. Activation of this pathway, often initiated by growth factors binding to their receptors, leads to the phosphorylation of Akt, which in turn phosphorylates and modulates a wide array of downstream effectors. One critical downstream target of Akt is the anti-apoptotic protein Bcl-2, whose upregulation promotes cell survival. A growing body of evidence from other cell systems indicates that Src can engage in complex crosstalk with the PI3K/Akt pathway, either positively or negatively regulating its activity. For instance, in some cancer models, Src contributes to the activation of this survival cascade [3], while in other contexts, such as in the mechanism of action of natural compounds like evodiamine, Src inhibition can lead to the suppression of proliferation and induction of apoptosis [4]. This context-dependent signaling underscores the complexity of Src function. Importantly, a direct link between Src and the regulation of the PI3K/Akt/Bcl-2 axis in human trophoblast cells remains unexplored. Establishing whether Src acts as a positive or negative regulator of this survival pathway in trophoblasts is essential to decipher its specific role in placentation.

To address this gap, the present study employs a loss-of-function approach using the immortalized human extravillous trophoblast cell line HTR-8/SVneo, a widely accepted and well-characterized in vitro model for studying trophoblast biology [1]. Through the application of specific small interfering RNA (siRNA) to knock down Src expression, we aimed to systematically evaluate the impact of its reduction on trophoblast cell function. The advantages of this RNA interference strategy include its high specificity, which enables a focused analysis of the target gene with minimized off-target effects, and its suitability for transient gene silencing to observe acute phenotypic changes without the potential confounding effects of long-term compensation seen in stable knockout models. By combining this genetic manipulation with a comprehensive suite of functional assays—including assessments of cell proliferation (CCK-8), migration (wound healing), invasion (Transwell with Matrigel), and apoptosis (flow cytometry and Western blot for cleaved caspases)—we can capture a holistic view of Src’s influence on trophoblast behavior.

The primary objective of this study, therefore, was to characterize the functional role of Src in controlling the proliferation, migration, invasion, and apoptosis of HTR-8/SVneo trophoblast cells. A secondary, and equally critical, aim was to delineate the underlying molecular mechanisms by investigating the effect of Src knockdown on the expression of key components of the PI3K/Akt/Bcl-2 signaling pathway. By probing whether Src modulates the expression of functionally relevant proteins, such as the p85 regulatory subunit of PI3K and the survival factor Bcl-2, we sought to define the intracellular signaling context through which Src exerts its effects. This integrative approach, linking a specific molecular perturbation to a wide range of cellular phenotypes and downstream signaling alterations, is designed to generate robust and comprehensive evidence regarding the role of Src in trophoblast biology. Ultimately, this work aims to provide foundational knowledge that could inform future therapeutic strategies for pregnancy complications characterized by trophoblast dysfunction, such as early pregnancy loss and preeclampsia.

## 2. Materials and Methods

### 2.1 Cells and Cell Culture

HTR8/SVneo cells were cultured in RPMI 1640 medium (Gibco, Cat. no. 21875-042) supplemented with 10% fetal bovine serum (FBS; Gibco, Cat. no. 10099141). The cells were maintained in a humidified incubator at 37°C with 5% CO_2_. Upon reaching 80%–90% confluency, cells were digested and passaged at a ratio of 1:2 using 0.25% Trypsin-EDTA (Gibco, Cat. no. 25200056) to ensure they remained in the logarithmic growth phase for subsequent experiments. JAR cells were cultured using the same protocol as HTR8/SVneo cells.

### 2.2 Small Interfering RNA (siRNA) Treatment

Cells were seeded into six-well plates and cultured for 24 h in complete medium prior to transfection. Transfection was performed using Lipofectamine 6000 (Beyotime, Cat. no. C0526) according to the manufacturer’s instructions. Cells were transfected with either control siRNA or SRC-specific siRNA oligonucleotides (Ribo, Cat. no. SIGS0007750-1). Following transfection, cells were cultured in complete medium and incubated at 37°C for 24 h and 48 h.

### 2.3 Western Blot Analysis

Total protein was extracted from cells using RIPA buffer (Beyotime, Cat. no. P0013B) following the manufacturer’s instructions. After centrifugation, the supernatant was collected, and protein concentration was quantified using a BCA assay kit (Zeta Life, Cat. no. ZT10012). Equal amounts of protein were separated by SDS-PAGE (Beyotime, Cat. no. P0015L) and transferred onto PVDF membranes (Beyotime, Cat. no. FFP33). Membranes were blocked with 5% non-fat milk in TBST for 1 h at room temperature and then incubated overnight at 4°C with primary antibodies (Beyotime, Cat. no. AF1966, AF0045, AF7299, AF1213, AF1264, AF5009, AT819). After three washes with TBST, membranes were incubated with HRP-conjugated secondary antibodies for 1 h at room temperature. Protein bands were visualized using an ECL detection system (Beyotime, Cat. no. P0018AM). Band intensities were normalized to an internal reference protein and quantified using ImageJ software.

### 2.4 RT-qPCR Analysis

Total RNA was extracted using a miRNA Isolation Kit (Tiangen, Cat. no. DP419) and reverse-transcribed into cDNA using a miRNA RT Kit (Tiangen, Cat. no. KR116-01) with miRNA-specific stem-loop primers to enhance specificity. Quantitative real-time PCR (qRT-PCR) was performed using SYBR Green qPCR Mix (Tiangen, Cat. no. RK145) and the aforementioned primers. Expression levels were normalized to U6 small nuclear RNA, and relative gene expression was calculated using the 2−ΔΔCt method.

### 2.5 Cell Counting Kit-8 (CCK-8) Assay

Cell proliferation was assessed using the Cell Counting Kit-8 (CCK-8) assay (Zeta Life, Cat. no. K009). HTR8 cells were seeded in 96-well plates at a density of 2,000 cells/well, transfected the following day, and cultured continuously for 48 h. CCK-8 reagent was added according to the manufacturer’s protocol. On days 0, 1, 2, 3, and 4, the culture medium was aspirated, and 100 μL of fresh complete medium containing 10 μL of CCK-8 reagent was added to each well. After incubation at 37°C for 1 h, absorbance at 450 nm (OD450) was measured to evaluate cell proliferation.

### 2.6 Cell Wound Scratch Assay

A wound healing assay was conducted to evaluate the migratory capacity of cells following SRC knockdown. Five parallel lines were marked on the bottom of 6 cm cell culture dishes. Transfected cells were seeded into the dishes and cultured until 90% confluency. A single linear scratch was made across the cell monolayer using a 10 μL pipette tip perpendicular to the pre-marked lines to ensure uniform scratch width across all samples. Cells were gently rinsed twice with PBS to remove detached cells and incubated in serum-free RPMI 1640 medium at 37°C with 5% CO_2_. Wound closure was monitored by imaging the scratched areas at 0, 6, 12, 24, and 48 h post-scratching.

### 2.7 Transwell Invasion Assay

Cell invasive capacity was assessed using a Transwell invasion assay. Briefly, 60 μL of Matrigel was coated onto the upper surface of each Transwell chamber (Corning, Cat. no. CLS3422) according to the manufacturer’s instructions, incubated at 37°C for 2 h, and excess liquid was removed. The chambers were hydrated with 100 μL of serum-free RPMI 1640 medium for 30 min. A suspension of 4 × 10^4^ HTR8 cells in 200 μL of serum-free RPMI 1640 medium was added to the upper chamber, while 500 μL of RPMI 1640 medium supplemented with 10% FBS was added to the lower chamber. After incubation at 37°C with 5% CO_2_ for 24 h, cells were stained with 0.5% crystal violet for 10 min. Uninvaded cells on the upper surface of the filter were wiped off with a cotton swab. Three random fields per chamber were imaged under a light microscope, and invaded cells were counted using ImageJ software.

### 2.8 Flow Cytometry Analysis

Apoptosis levels in HTR8 cells were detected using an Annexin V-FITC/PI Apoptosis Detection Kit (Yeasen, Cat. no. 40302). Two days post-transfection, HTR8 cells were harvested using EDTA-free trypsin, washed with pre-cooled PBS, and processed according to the kit instructions. Approximately 1 × 10^5^ to 5 × 10^5^ cells were resuspended in 100 μL of binding buffer in a centrifuge tube, followed by the addition of 5 μL Annexin V-FITC and 10 μL PI staining solution. After incubation in the dark for 10 min, 400 μL of binding buffer was added, and samples were analyzed by flow cytometry.

## 3. Result

### 3.1 Validation of SRC Knockdown Efficiency

To investigate the functional role of SRC, we performed siRNA-mediated knockdown to suppress endogenous SRC expression in HTR8 cells. HTR8 cells were transiently transfected with SRC siRNA (si-SRC), while cells transfected with non-targeting scrambled siRNA served as the negative control (NC), and cells treated with transfection reagent alone served as the blank control (Li group). RT-qPCR analysis revealed a significant decrease in SRC mRNA expression in si-SRC–transfected cells compared with control cells (Fig.1a). Western blot analysis further confirmed that SRC protein levels were markedly downregulated in HTR8/SVneo cells following SRC silencing (Fig.1b), demonstrating successful knockdown of SRC at both the mRNA and protein levels.

**Fig. 1.**
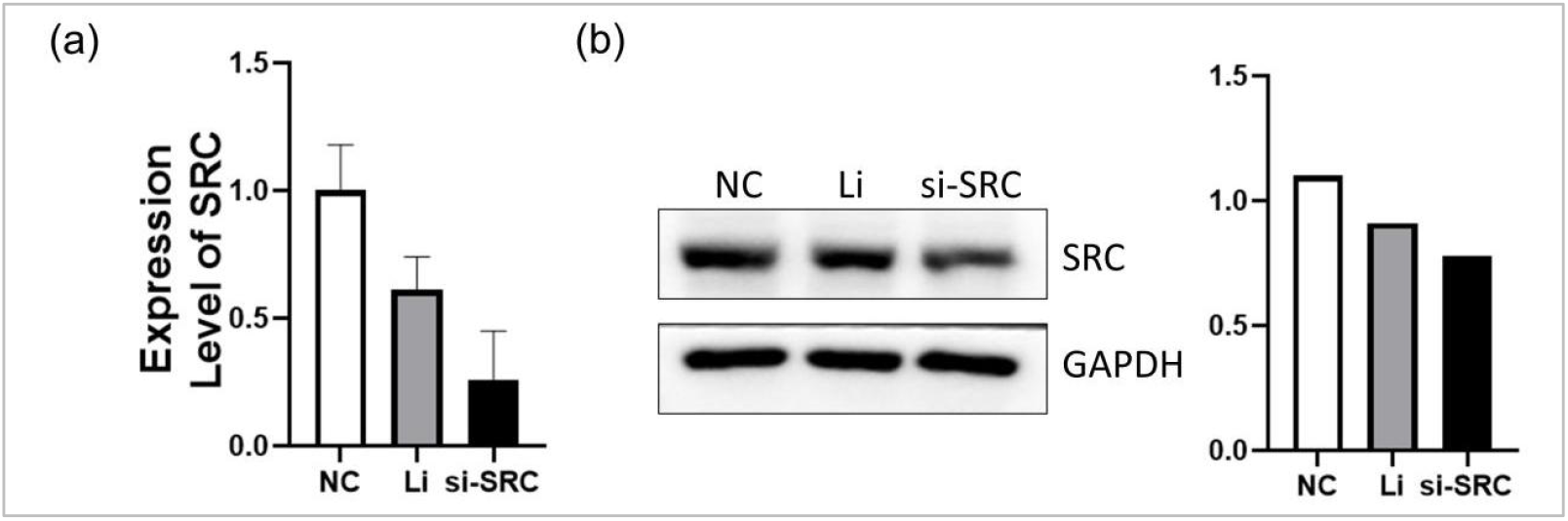
Validation of SRC knockdown efficiency in HTR8 cells. (a) Relative SRC mRNA expression detected by qPCR; (b) Relative SRC protein expression detected by Western blot.

### 3.2 SRC Knockdown Inhibits Trophoblast Cell Proliferation

Consistently, CCK-8 assays revealed significantly lower absorbance values in the si-SRC group, demonstrating impaired proliferative capacity in HTR8 cells upon SRC downregulation (Fig.2). Collectively, these data suggest that SRC depletion markedly attenuates the proliferative ability of trophoblast cells.

**Fig. 2.**
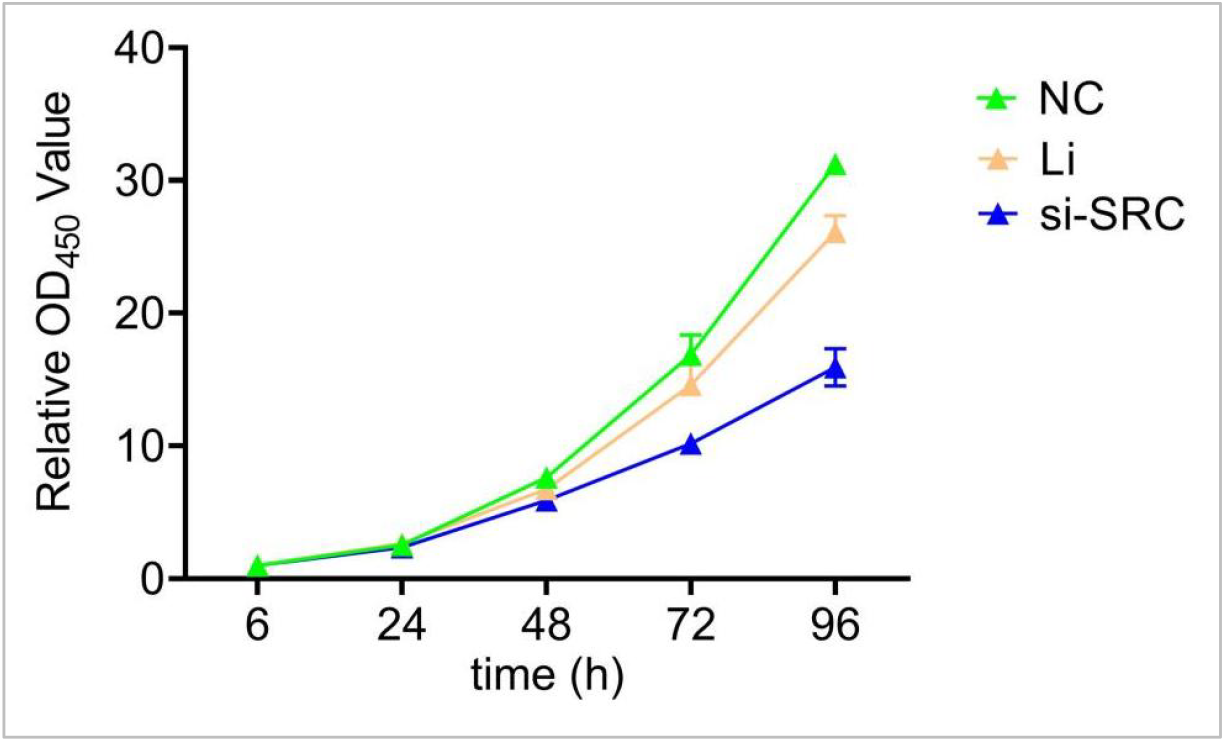
SRC knockdown inhibits trophoblast cell proliferation. Proliferation of HTR8 cells detected by CCK-8 assay (OD450 values).

### 3.3 SRC Knockdown Impairs Cell Migration and Invasion

Cell migration and invasion are essential cellular processes tightly regulated in physiological contexts such as embryonic development and tissue homeostasis; their dysregulation is closely associated with various pathological conditions, including abnormal trophoblast function. To further clarify the regulatory role of SRC in HTR8 cell migration and invasion, we employed scratch and Transwell assays to assess functional changes following SRC knockdown. Scratch assay results showed that the wound healing rate of HTR8 cells in the si-SRC group was significantly lower than that in the NC and Li groups, indicating that SRC knockdown significantly impaired the migratory capacity of HTR8 cells (Fig.3). Similarly, Transwell invasion assay results demonstrated that the number of invasive HTR8 cells in the si-SRC group was markedly reduced compared with the NC and Li groups (Fig.4). No significant differences in wound healing rate or the number of invasive cells were observed between the NC and Li groups, suggesting that the transfection reagent itself had no effect on HTR8 cell migration and invasion.

**Fig. 3.**
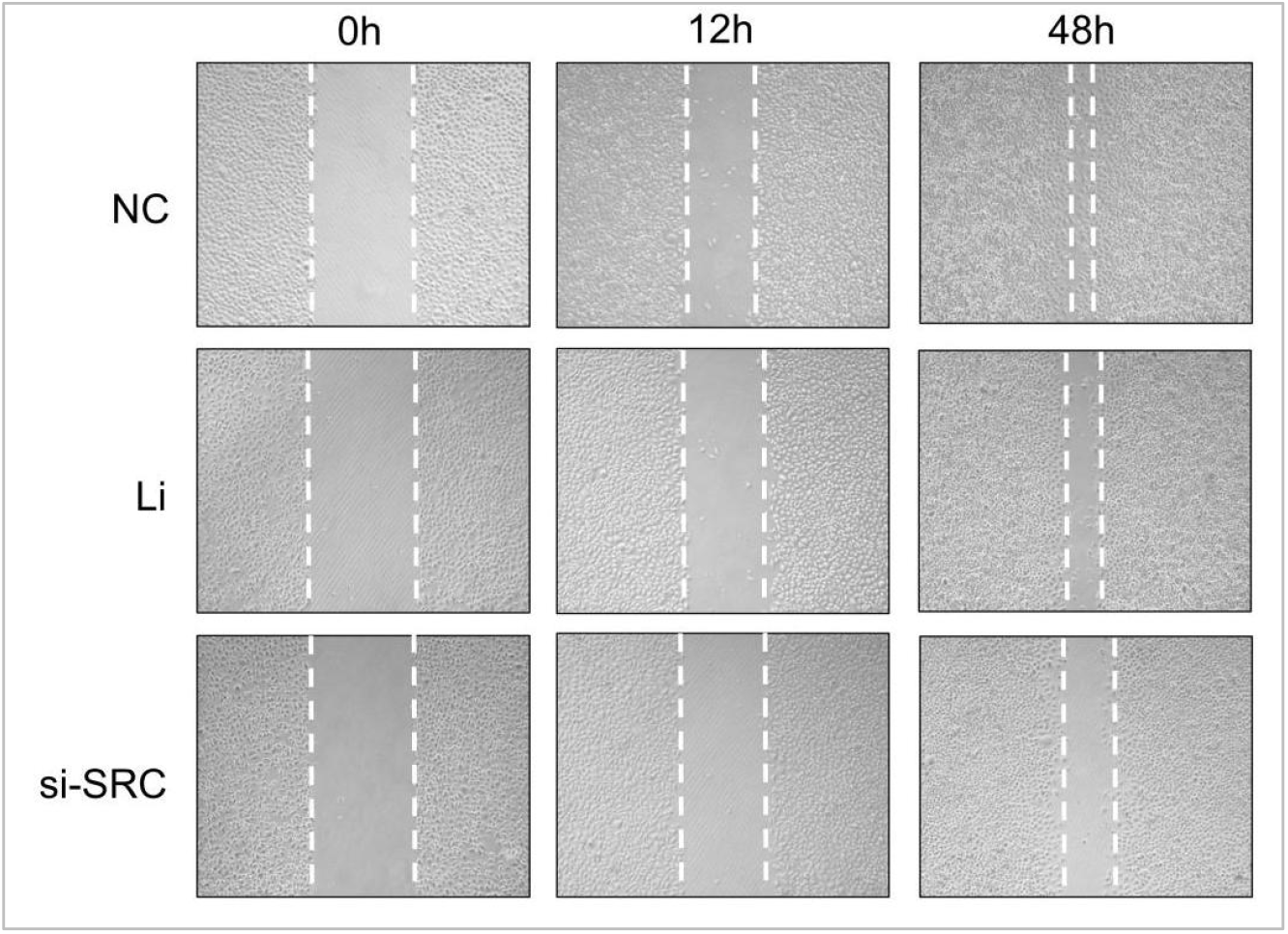
Wound healing rate of HTR8/SVneo cells detected by scratch assay.

**Fig. 4.**
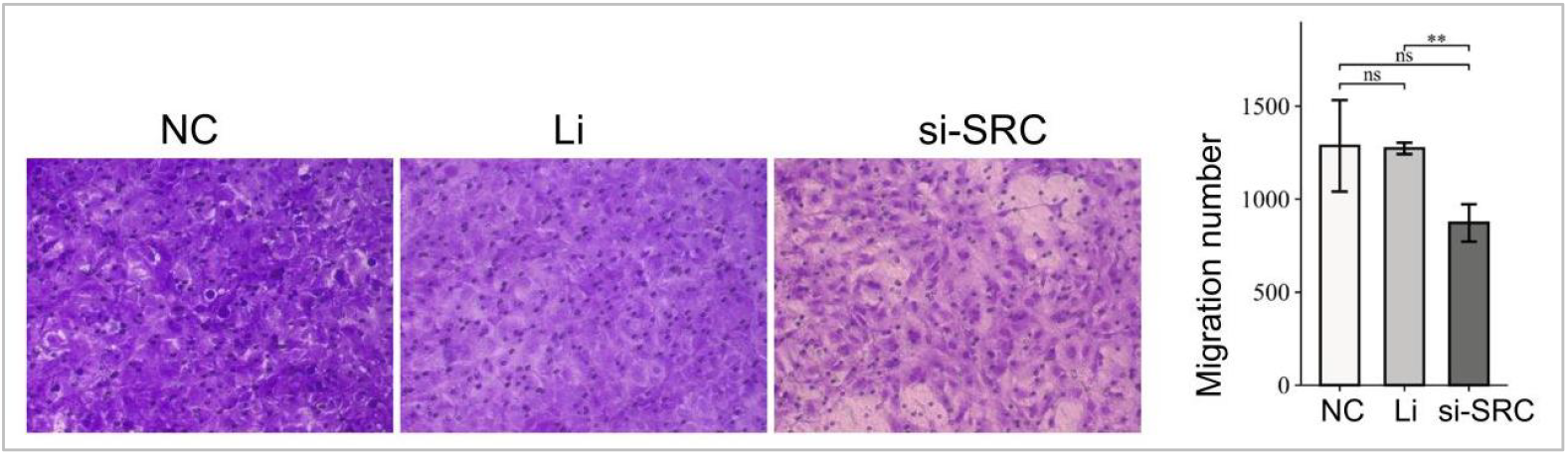
Number of invasive HTR8/SVneo cells detected by Transwell assay.

### 3.4 SRC Depletion Induces Apoptosis via the Caspase-Dependent Pathway

To determine whether SRC depletion induces apoptosis in trophoblast cells, we performed Annexin V-FITC/PI double staining followed by flow cytometry in HTR8 cells. Apoptotic cells were defined as those located in quadrant Q3 (early apoptosis) and quadrant Q2 (late apoptosis). Representative flow cytometry dot plots revealed a marked increase in the proportion of apoptotic cells in the si-SRC group compared with both the NC and Li groups. Quantitative analysis confirmed that the apoptosis rate in the si-SRC group was significantly higher than that in the NC and Li groups (Fig.5). To further validate the pro-apoptotic effect of SRC knockdown, we performed Western blot analysis to detect the expression levels of apoptosis-related proteins Caspase-3 and Caspase-9. The results showed that protein levels of Caspase-3 and Caspase-9 were significantly upregulated in the si-SRC group compared with the NC and Li groups. These Western blot results were consistent with the flow cytometry data, further confirming that SRC knockdown induces apoptosis in HTR8 cells by activating the Caspase-dependent apoptotic pathway (Fig.6). Collectively, these findings demonstrate that SRC knockdown promotes apoptosis in trophoblast cells, underscoring its critical role in maintaining trophoblast cell survival.

**Fig. 5.**
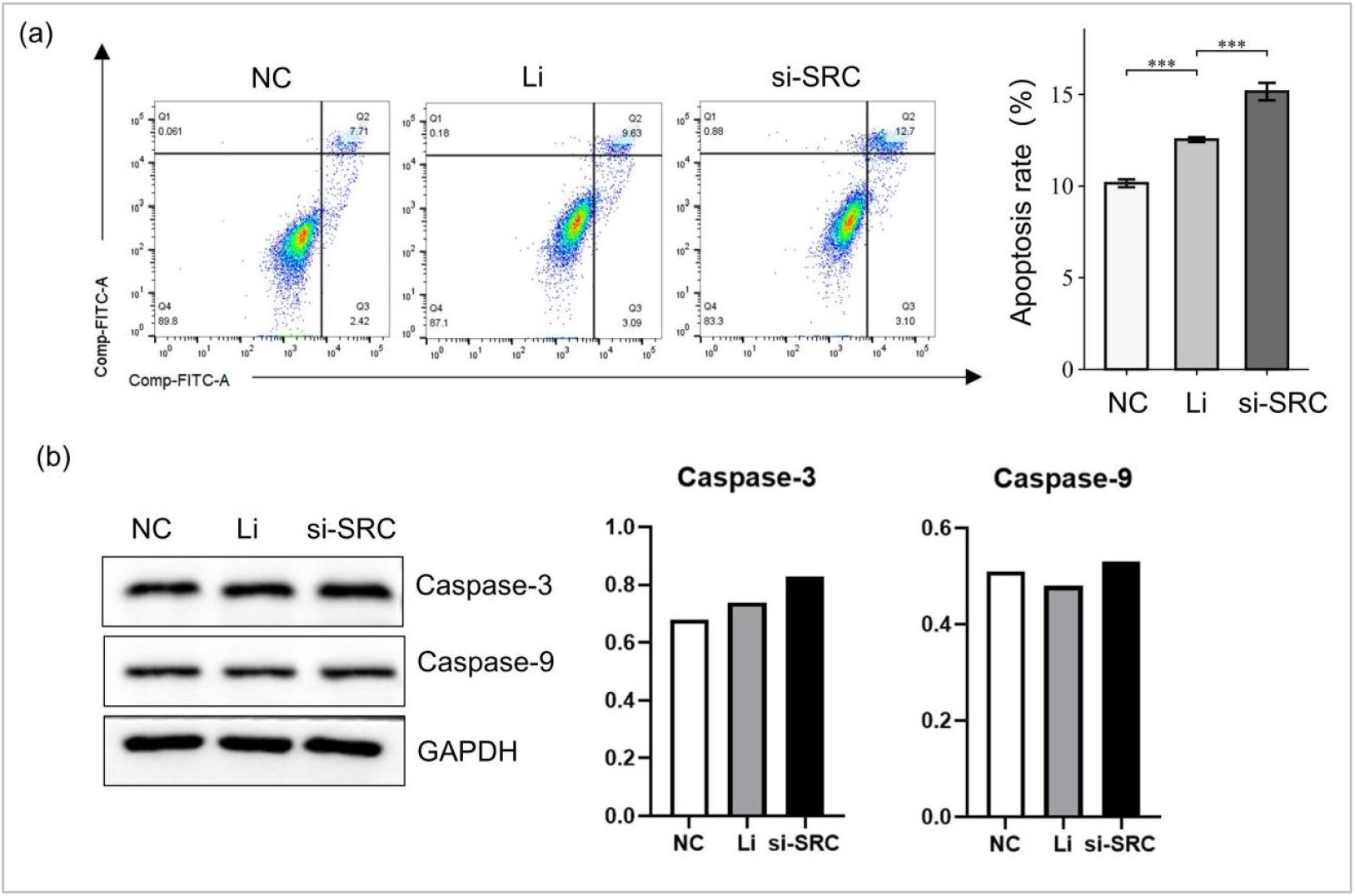
SRC knockdown enhances apoptosis of HTR8/SVneo cells detected by flow cytometry.

**Fig. 6.**
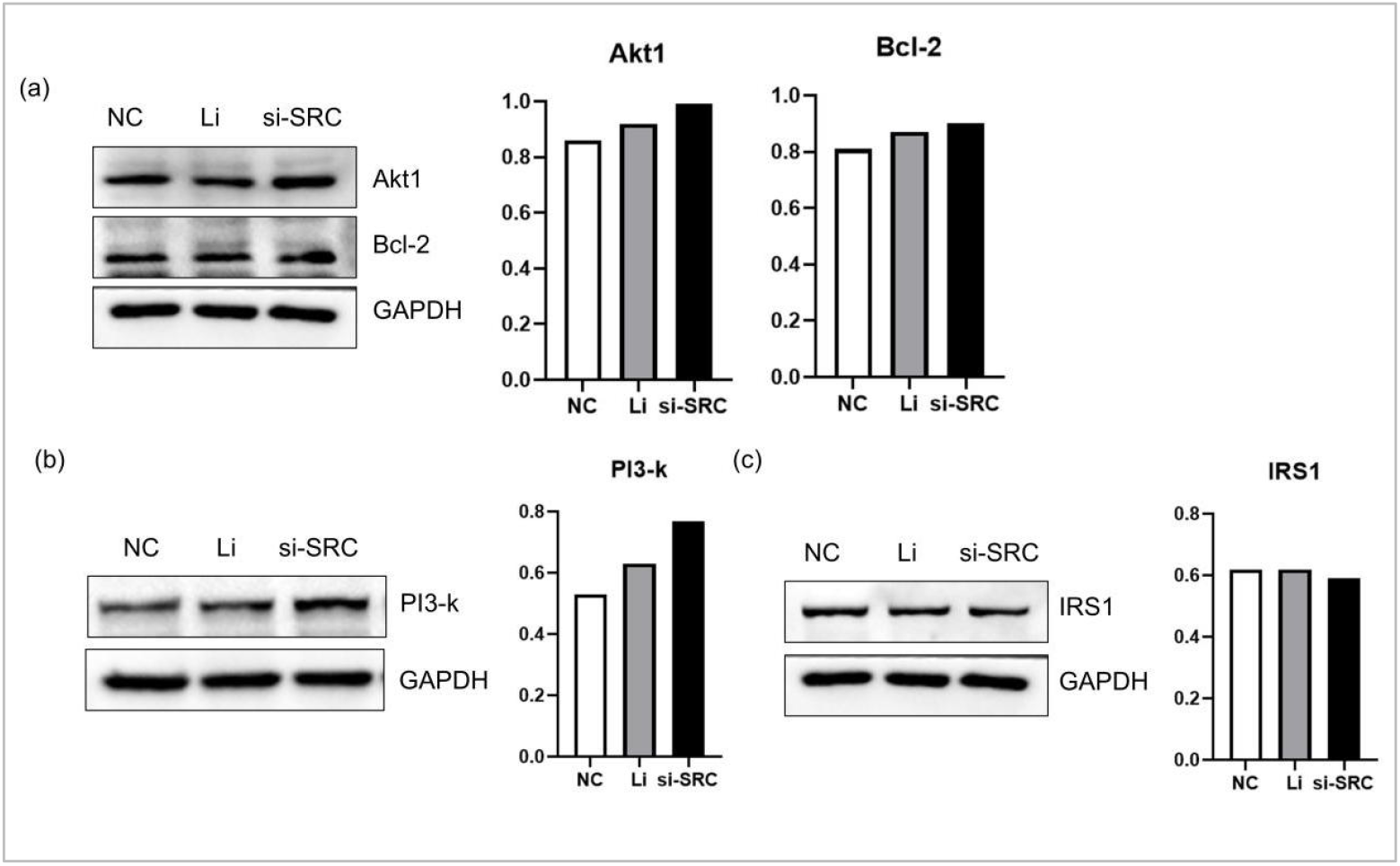
SRC knockdown activates the PI3K/Akt/Bcl-2 signaling pathway in HTR8 cells.

### 3.5 SRC Knockdown Activates the PI3K/Akt/Bcl-2 Signaling Pathway

The PI3K/Akt signaling axis is a central regulator of cell survival, proliferation, and apoptosis in trophoblasts, mediating downstream anti-apoptotic gene expression and promoting cell viability. To elucidate the molecular mechanism by which SRC modulates HTR8 trophoblast cell function, we performed Western blot analysis to examine the expression of core components in the PI3K/Akt signaling pathway, including PI3K, Akt1, Bcl-2, and IRS1, following SRC knockdown (si-SRC). NC and Li groups were included to account for non-specific effects of transfection. As depicted in Fig.6, SRC knockdown led to a marked upregulation of PI3K protein expression, accompanied by elevated levels of its downstream kinase Akt1 and the anti-apoptotic effector Bcl-2. In contrast, the expression of IRS1, an upstream adaptor molecule linking receptor tyrosine kinases to PI3K activation, remained largely unchanged relative to control groups. These findings suggest that SRC acts as a negative regulator of the PI3K/Akt/Bcl-2 survival axis in HTR8 cells. Given that IRS1 expression was unaltered, the observed activation of PI3K likely occurs through an IRS1-independent mechanism, possibly via direct SRC-mediated phosphorylation of PI3K regulatory subunits or modulation of PI3K stability. The consequent increase in Akt1 activity further promotes the upregulation of Bcl-2, thereby enhancing cell survival and inhibiting apoptosis. Collectively, these findings demonstrate that SRC knockdown activates the PI3K/Akt/Bcl-2 pathway, ultimately leading to alterations in HTR8 cell function.

## 4. Discussion

Trophoblast cell function is fundamental to successful embryo implantation and placentation, and its dysregulation is a core pathological mechanism underlying pregnancy complications such as preeclampsia, miscarriage, and fetal growth restriction [1, 5]. The extravillous trophoblast must precisely coordinate proliferation, migration, and invasion to remodel the maternal spiral arteries, while aberrant apoptosis disrupts placental homeostasis [6]. Although the proto-oncogene Src (SRC) is a well-established signal transducer mediating cell survival, migration, and cytoskeletal dynamics in multiple cell types, its specific roles in trophoblast biology remain incompletely defined [2,7]. Specifically, whether SRC modulates trophoblast function via the PI3K/Akt/Bcl-2 survival axis, a pathway classically associated with anti-apoptotic signaling, is unknown [8].

This study systematically investigated the functional role of SRC in the HTR-8/SVneo extravillous trophoblast cell line using siRNA-mediated knockdown combined with comprehensive phenotypic and molecular analyses. Our results demonstrate that SRC silencing significantly impairs trophoblast proliferation, migration, and invasion, while concurrently promoting caspase-dependent apoptosis. Paradoxically, this suppression of trophoblast function was accompanied by the upregulation of PI3K, Akt1, and Bcl-2 protein expression, indicating that SRC normally exerts a negative regulatory effect on the PI3K/Akt/Bcl-2 signaling cascade. These findings reveal a novel dual role for SRC: it acts as a positive regulator of trophoblast motility and survival, while simultaneously restraining a pro-survival pathway. In the following discussion, we will interpret these key findings in the context of existing literature, explore their pathophysiological implications for pregnancy disorders, and examine the mechanistic paradox arising from the concurrent activation of a survival pathway despite the loss of cellular function.

The initial confirmation of efficient SRC knockdown at both the mRNA and protein levels in HTR-8/SVneo cells provides a robust foundation for subsequent functional analyses. This specificity is critical, as off-target effects are a primary concern with siRNA-based approaches; the inclusion of both a negative control (NC) and a lipofectamine-only (Li) group effectively demonstrates that the observed phenotypes are directly attributable to SRC depletion rather than transfection stress or off-target gene silencing [7]. The choice of the HTR-8/SVneo cell line is particularly relevant, as it is a well-established model for studying human extravillous trophoblast function; indeed, genome-scale studies have identified an SRC-dependent transcriptome in these cells that encodes proteins essential for placental tissue development [1]. While CRISPR-Cas9 offers the advantage of stable gene knockout, the transient nature of siRNA-mediated knockdown allows for the assessment of acute SRC loss, which can mimic rapid regulatory changes that occur in the placental environment. Furthermore, the dose-dependent nature of this knockdown suggests that even partial reduction of SRC protein is sufficient to trigger significant downstream signaling alterations, a point that underscores the gene’s potent regulatory role in trophoblast biology.

Building upon the validated knockdown model, the significant suppression of HTR-8/SVneo cell proliferation highlights SRC as a critical positive regulator of trophoblast growth. This finding aligns with the broader oncogenic role of SRC in driving cell cycle progression, and it resonates with observations in other gestational tissues, where signaling through the SRC-related kinase Shp2 is crucial for maintaining trophoblast cell cycle progression via suppression of the p53-p21 axis [2]. The observed proliferative deficit suggests that SRC may be integrated into a network of growth-promoting pathways, for instance, by acting downstream of receptor tyrosine kinases like EGFR, which has been directly shown to activate Rap1 and subsequently Src signaling to drive trophoblast invasion [9]. The functional consequences of SRC loss extend beyond simple growth arrest; the concurrent inhibition of migration and invasion reveals a pleiotropic role for SRC in trophoblast cell dynamics. This is highly consistent with previous work demonstrating that suppression of the FAK/SRC/PI3K/AKT pathway is a central mechanism by which environmental toxins impair trophoblast motility [10]. The ability of SRC to orchestrate both migration and invasion is likely mediated through its well-established role in regulating focal adhesion dynamics and cytoskeletal remodeling, potentially via downstream effectors such as the WAVE2-ARP2/3 complex, which has been shown to be crucial for actin reorganization and migration in these same HTR-8/SVneo cells [6].

In stark contrast to the inhibition of proliferative and migratory functions, SRC knockdown unexpectedly led to a pronounced increase in apoptosis, as evidenced by elevated rates of Annexin V positivity and the upregulation of Caspase-3 and -9. This dual phenotype—impaired cell function yet increased cell death—is a hallmark of SRC’s multifaceted role, acting not only as a promoter of cell motility but also as a key survival factor in trophoblasts. This pro-survival role is reinforced by studies showing that the SRC/LDHA pathway in trophoblast-derived lactic acid signaling is essential for early pregnancy maintenance, and that disruptions to this pathway are linked to recurrent pregnancy loss [11]. The activation of the initiator caspase-9 strongly implicates the intrinsic, mitochondrial-mediated apoptotic pathway. This finding is mechanistically significant, as SRC is known to exert anti-apoptotic effects by phosphorylating and inactivating pro-apoptotic factors such as Bad, a protein that directly regulates the mitochondrial permeability transition pore.

The most compelling and paradoxical finding of this study is the simultaneous activation of the PI3K/Akt/Bcl-2 survival pathway following SRC knockdown. While the upregulation of PI3K, p-Akt, and Bcl-2 is traditionally associated with enhanced cell survival and proliferation, these outcomes were not observed; indeed, the cells exhibited suppressed growth and increased death[12,13]. This suggests a complex feedback mechanism where the cell attempts to compensate for SRC loss by hyperactivating an alternate survival axis. A plausible explanation for this phenomenon is the removal of a negative regulatory influence exerted by SRC on the PI3K pathway. For instance, SRC can activate the phosphatase PTEN, a major negative regulator of PI3K signaling; thus, SRC knockdown could relieve PTEN activity, leading to an increase in PI3K product levels [14]. The lack of change in IRS1 expression further suggests that this compensatory activation occurs at a level downstream of direct receptor engagement. This disconnect between a pro-survival signaling output and a pro-apoptotic functional outcome highlights the dominance of SRC-regulated pathways—perhaps MAPK/ERK or STAT3—that are indispensable for trophoblast function and whose absence cannot be compensated for by PI3K/Akt activation alone [15]. This nuanced interaction provides a new perspective on how the SRC-PI3K/Akt axis is wired in placental cells.

Several limitations of this study should be acknowledged. First, all experiments were conducted exclusively using the immortalized HTR8/SVneo trophoblast cell line, which, while widely used, may not fully recapitulate the biological characteristics of primary trophoblasts or the complex in vivo microenvironment. The absence of validation in primary cells or trophoblast-specific SRC knockout animal models limits the generalizability of our findings to physiological and pathological pregnancy conditions. Second, the mechanistic exploration remains incomplete. Although we observed that SRC knockdown paradoxically activated the PI3K/Akt/Bcl-2 survival pathway while inhibiting proliferation and migration, the study did not investigate other critical signaling cascades (e.g., MAPK/ERK, STAT3) that may compensate or interact. Furthermore, the precise molecular mechanism by which SRC negatively regulates PI3K/Akt—such as potential involvement of PTEN or direct phosphorylation of PI3K regulatory subunits—was not elucidated. The single time-point analysis (48 hours post-knockdown) also fails to capture temporal dynamics of pathway activation.

In summary, this study provides compelling evidence that SRC is a critical positive regulator of trophoblast cell proliferation, migration, and invasion while exerting anti-apoptotic effects in HTR8/SVneo cells. Mechanistically, SRC knockdown activates the PI3K/Akt/Bcl-2 pathway, likely as a compensatory response, but this is insufficient to overcome the functional deficits caused by SRC loss. These findings reveal a novel regulatory role of SRC in trophoblast biology and offer potential insights into the pathogenesis of pregnancy disorders such as preeclampsia and miscarriage. Future research should focus on validating these results in primary trophoblast cells and in vivo models, and on dissecting the upstream molecular events that link SRC to PI3K/Akt inhibition. Targeting SRC may represent a promising therapeutic strategy for modulating trophoblast function in pregnancy-related diseases.

## Notes

**Funding sources** This research was supported by Zhejiang Provincial Natural Science Foundation of China under Grant No.LTGY23H040004, No.LTGY23H040005 , No.LTGY24H040006, No.LTGY24H040005 , No.LKLY26H040001 and the Health Commission of Zhejiang Province, China (WKJ-ZJ-2449, 2025HY1308, 2025A14016, 2024A14004) , the Science Technology Department of Shaoxing, China (2022A14006, 2025A14003, 2025A14006),the Health Commission of Shaoxing, China (2025A14016 , 2025SKY044 , 2025SKY047 , 2024SKY003, 2022KY036, 2022KY038, 2025SKY045 , 2023SKY047).Key Laboratory of Reproductive Health of Shaoxing City, Shaoxing Maternity and Child Health Care Hospital, Shaoxing, Zhejiang 312000, China(2023SSY004, 2023SSY007). Zhejiang University Shaoxing Research Institute - Shaoxing Maternal and Child Health Care Hospital Joint Research Grant for Life and Health (2025SMJK004).

### Competing Interest Statement

The authors have declared no competing interest.

